# Transient Receptor Potential Ankyrin1 channel is endogenously expressed in T cells and regulates immune functions

**DOI:** 10.1101/621854

**Authors:** Subhransu Sekhar Sahoo, Rakesh Kumar Majhi, Ankit Tiwari, Tusar Acharya, P Sanjai Kumar, Somdatta Saha, Chandan Goswami, Subhasis Chattopadhyay

## Abstract

Transient Receptor Potential channel subfamily A member 1 (TRPA1) is a non selective cationic channel, identified initially as a cold sensory receptor. TRPA1 responds to diverse exogenous and endogenous stimuli associated with pain and inflammation. However, the role of TRPA1 towards T cell responses remains scanty. In this work, we explored the endogenous expression of TRPA1 in T cells. By RT-PCR we confirmed the expression of TRPA1 in T cell at RNA level. Using confocal microscopy as well as flow cytometry, we demonstrated that TRPA1 is endogenously expressed in primary murine splenic T cells as well as in primary human T cells. The endogenous expression of TRPA1 is confirmed by using another antibody. TRPA1 was primarily located at the cell surface. TRPA1-specific activator namely AITC increases intracellular Ca^2+^-levels while two different inhibitors namely A-967079 as well as HC-030031 reduce intracellular Ca^2+^-levels in T cells. Such Ca^2+^-influx can also be influenced by chelation of intracellular Ca^2+^ as well as extracellular Ca^2+^. TRPA1 expression was found to be increased during αCD3/αCD28 (TCR) or ConA driven stimulation in T cells. TRPA1-specific inhibitor treatment prevented induction of CD25, CD69 in ConA/TCR stimulated T cells and secretion of cytokines like TNF, IFN-γ and IL-2 suggesting that endogenous activity of TRPA1 may be involved in T cell activation. Collectively these results may have implication in T cell-mediated responses and possible role of TRPA1 in immunological disorders.

## Introduction

Transient receptor potential cation channel subfamily A member 1 (TRPA1) is the only member of the mammalian ‘ankyrin’ type subfamily of TRP channels [1]. TRPA1 is a Ca^2+^-permeable non-selective cation channel that is expressed in a set of nociceptive/thermo-receptive neurons that can detect noxious cold temperature below 17°C. It is often co-expressed with the heat-sensitive channel TRPV1 rather than with the cold-sensitive channel TRPM8. Initial reports suggested that the large influx of Ca^2+^ ions at lower temperatures (such as at 10°C) is primarily due to the direct activation of this channel by cold stimuli [2, 3]. Apart from low temperature as a stimulus, TRPA1 also acts as a nociceptive receptor that detects noxious chemicals that can cause tissue damage. TRPA1 also acts as a mediator of inflammatory pain generated by either noxious cold or chemical irritants [4, 5]. Ca^2+^ channels are an integral part of T cell activation, differentiation, and formation of an immunological synapse between mature CD4^+^ T cells and Antigen Presenting Cells (APC), and even exocytosis of vesicles in cytotoxic T cells. It also influences cytokine release patterns which in turn affect T cell functions like the development of anergy, maturation, and differentiation of naïve T cells into Th1 andTh2 cells [6]. In agreement with these reports, abnormality in Ca^2+^-signalling results in several immunological disorders such as SCID, Wiskott–Aldrich syndrome (WAS)[6]. T cell motility is also regulated by Ca^2+^ in consortium with Protein Kinase C which regulates the rearrangement of actin cytoskeleton in them. Activation of several transcription factors such as NFAT, NF-κB, calmodulin-dependent kinase, JNK, and others are known to be involved in T cell regulation and require Ca^2+^ ions [7]. Moreover, recently a strategic usage of cation channel blockers towards T cell activation, cytokine secretion, and proliferation has been demonstrated [8]. Transient Receptor Potential (TRP) channels act as Ca^2+^-permeable channels and thereby regulate Ca^2+^-homeostasis. So far few members of the TRP family members have been identified by us and other groups that are endogenously expressed in T cells and regulate T cell functions [9, 10] and us. In this work, we have probed for the expression and localization of the TRPA1 channel in T cells and explored its functional importance as well.

Endogenous inflammatory agents such as Reactive Oxygen Species (ROS) induce the production of 4-hydroxy-2-nonenal (HNE) that directly activates TRPA1 [9]. Nitrated fatty acids and prostaglandins like 15d-PGJ2 are released at the site of inflammation, and such compounds can also directly activate TRPA1 [10, 11]. It is also known that lymphoma and inflammation associated itch is mediated by a Th2 cell-derived cytokine, interleukin-31 (IL-31), that directly interacts with its receptor IL-31RA located on TRPV1^+^/TRPA1^+^ sensory nerves in skin [12]. These compounds can also play a role in T cell activation. In addition, LPS (a noxious by-product of gram-negative bacteria) activates TRPA1 via a TLR-4 independent mechanism and thereby generates a rapid nociceptive response and neurogenic inflammation [13]. In this work, we have probed for the expression, localization of TRPA1 channel in T cells and also characterized the function of TRPA1 towards T cell activation.

## Materials and Methods

### Reagents

The TRPA1 channel modulatory drugs Allyl isothiocyanate (AITC, activator), A-967079 (inhibitor) and HC-030031 (another inhibitor) were obtained from Sigma-Aldrich (St Louis, MO, USA). ConA was purchased from Himedia (Mumbai, India). Rabbit polyclonal antibody against the 1st extracellular loop (747-760aa) of hTRPA1 and it’s specific blocking peptide (NSTGIINETSDHSE) were purchased from Alomone Laboratories (Jerusalem, Israel). The other rabbit polyclonal antibody against the N-terminus of TRPA1 was procured from Novus Biologicals. The Ca^2+^-sensitive dye Fluo-4AM was procured from Molecular Probes (Eugene, OR, USA). Calcium chelating agents BAPTA-AM and EGTA were procured from Sigma (St Louis, MO, USA). Trizol was purchased from Life Technologies (Carlsbad, CA, USA). Verso cDNA synthesis kit was obtained from ThermoScientific (Waltham, MA, USA). SYBR Green for RT-PCR was procured from Life Technologies (Carlsbad, CA, USA). The RT-PCR Primers for TRPA1 and GAPDH were obtained from IDT (Coralville, IO, USA). Anti-mouse CD25PE, CD69PE, and anti-human CD3-PE as well as functional grade (Azide-free) anti-CD3 and anti CD28 mAbs were obtained from BD Biosciences (San Jose, CA, USA). The CD90.2 APC antibody was from Tonbo Biosciences (San Diego, CA, USA). The sequences of the primers used in the study are mentioned below:

TRPA1 Forward: 5’-GTC CAG GGC GTT GTC TAT CG -3’

TRPA1 Reverse: 5’-CGT GAT GCA GAG GAC AGA AT -3’

GAPDH Forward: 5’-CCG CAT CTT CTT GTG CAG TG-3’

GAPDH Reverse: 5’-CCC AAT ACG GCC AAA TCC GT-3’

### Isolation and culture of T cells

T cells were isolated and cultured as described previously [14]. Briefly, murine spleen cells were obtained from 6 to 8-week-old BALB/c mice as per the approval of the Institutional Animal Ethics Committee (IAEC protocol no. NISER/SBS/IAEC/AH-39). Single cell suspension was made by passing the suspended splenocytes through a 70 µm cell strainer. T cells were purified from the non-adherent splenocyte population by using BD IMag™ Mouse T Lymphocyte Enrichment Set – DM according to the manufacturer’s instructions. The isolated cells were cultured in a 24 well polystyrene cell culture plate (3.5 × 10^6^ cells/well) with RPMI (PAN Biotech, Aidenbach, Germany) supplemented with 10% FBS (PAN Biotech). The percentage purity of the purified T cells was above 95% in each case. All the experiments (assays) were performed about 36 hours after plating the cells as most of the primary T cells were found to be activated during 36–48 hours after ConA or TCR treatment (data not shown). Primary murine T cells were activated with either a combination of plate-bound α-CD3 (2µg/mL) and soluble α-CD28 (2µg/mL) or with ConA (4µg/mL) alone for 36 hours before experiments. Similarly, human PBMC derived T cells were purified by Human T cell isolation kit from Invitrogen (Invitrogen Dynal AS, Oslo, Norway) according to manufacturer’s instruction. Treatment of T cells was carried out with selective TRPA1 activator AITC (100µM) and inhibitor A-967079 (100µM) with or without ConA or anti-CD3/CD28.

### RNA isolation and qPCR

Approximately 3-5 × 10^6^ T cells were used for RNA isolation. For positive control, spinal cord tissue from mice was used. RNA isolation was done using TRIzol reagent according to the manufacturer’s instructions. Nanodrop readings were taken and 1 μg of RNA was converted to cDNA using verso cDNA synthesis kit as per the mentioned protocol. RTPCR reactions were performed for TRPA1 and GAPDH in ABI7500 system (Applied Biosystems, Foster city, CA, USA) using 2X SYBR Green mix following the reaction gene expression was visualized in 1 % agarose gel.

### Flow cytometry

Flow cytometry analysis of T cells was performed as described previously [14]. For probing for TRPA1 expression, cells were stained with the TRPA1-specific antibody mentioned before and subsequently flow cytometric analysis was performed as described previously [14, 15]. For evaluating the profile of immune markers, mouse T cells were incubated with anti-CD25 PE, CD69 PE and CD90.2 APC mAbs dissolved in FACS buffer (1 X PBS, 1% BSA and 0.05% sodium azide) for 30 min on ice and then washed further. Stained cells were washed twice with the same FACS buffer before the line-gated acquisition of around 10,000 cells. Human PBMC derived T cells were stained with anti-human CD3 antibody and were acquired with FACS Calibur (BD Biosciences). Data were analyzed using CELL QUEST PRO software (BD Biosciences). The percentages of cells expressing the markers are represented in dot-plots while the MFI values represent the expression levels of the markers per cell.

### Immunofluorescence analysis and microscopy

Immunofluorescence analysis of T cells was performed as described previously [14]. For immunocytochemical analysis, immediately after harvesting, T cells were diluted in PBS and fixed with paraformaldehyde (final concentration 2%). After fixing the cells with paraformaldehyde, immunostaining was done by two procedures: in first case cell permeabilization was not performed as the antibody detecting TRPA1 recognizes an extracellular region of TRPA1. In other cases, the cells were permeabilized with 0.1% Triton X-100 in PBS (5 min). Subsequently, the cells were blocked with 5% BSA for 1 hour. The primary antibodies were used at 1: 400 dilutions. In some experiments, blocking peptides were used to confirm the specificity of the immunoreactivity. The ratio of blocking peptides with specific antibody was 1: 1 (in concentration). Another rabbit polyclonal antibody (procured from Novus Biologicals) detecting epitope present in the N-terminal cytoplasmic domain of TRPA1 was used (1:1000 dilution) in some experiments to confirm the endogenous expression of TRPA1 in T cells. All primary antibodies were incubated overnight at 4°C in PBST buffer (PBS supplemented with 0.1% Tween-20). AlexaFluor-488 labeled anti-rabbit antibody (Molecular Probes) was used as secondary antibodies and was used at 1:1000 dilutions. All images were acquired on a confocal laser scanning microscope (LSM-780, Zeiss) with a 63 X objective and analyzed with the Zeiss LSM image examiner software and Adobe Photoshop.

### Ca^2+^-imaging

Ca^2+^ imaging of primary murine splenic T cells was performed as described previously with minor modifications [14, 16]. In brief, primary murine splenic T cells were loaded with Ca^2+^-sensitive dye (Fluo-4 AM, 2 µM for 30 min). Ca^2+^-chelation experiment was performed by treating the cells with 5 µM BAPTA-AM for 1 hour and extracellular Ca^2+^ was chelated with 5 µM EGTA for 1 hour. The cell suspension was added to the live cell chamber for Ca^2+^ imaging and images were acquired in every 5-sec intervals. The cells were stimulated with specific agonists alone or in a combination of agonists and antagonists as described. Fluo-4 AM signal was acquired using a Zeiss microscope (LSM 780) & Olympus microscope (FV3000) and with the same settings. The images were analyzed using LSM software & Fiji and intensities specific for Ca^2+^-loaded Fluo-4 are represented in artificial rainbow color with a pseudo scale (red indicating the highest level of Ca^2+^ and blue indicating the lowest levels of Ca^2+^). For quantification of the changes in the intracellular Ca^2+^-levels, fluorescence intensity of T cells present in view field was measured before and just after adding the drugs. Such values were plotted for relative changes.

### ELISA

Supernatants from the respective experiments were collected and stored at −80°C. ELISA for different T cell effector cytokines TNF, IL-2 and IFNγ were performed using BD Biosciences Sandwich ELISA kits (San Jose, CA, USA) as per the manufacturer’s instructions. The readings were acquired using a microplate reader (Bio-Rad iMARK) at 450 nm.

### Statistical tests

The primary data were imported into R software for statistical analysis. The ANOVA test was done for each data set to check the reliability and significance. The P value of < 0.05 was considered statistically significant. Data presented here are representative of three independent experiments. The significance values are as follows: *** = P between 0 and 0.001; ** = P between 0.001 and 0.01; * = P between 0.01 and 0.05; ns = P above 0.05.

## Results

### TRPA1 is expressed endogenously in primary murine and human T cells

Expression of TRPA1 in mRNA level in T cell is confirmed by RT-PCR (Fig. 1a). The surface expressions of specific ion channels are critical for signaling events. We used a specific antibody (Almone lab) for which the epitope is present at the extracellular loop-1 of TRPA1 (i.e., present outside the cell surface). Therefore this antibody allowed us to probe for the surface expression of TRPA1 (in un-permeabilized cells) and as well as total TRPA1 expression (in Triton X-100-permeabilized cells). We probed the surface expression of TRPA1 in unpermeabilized cells. This antibody detects TRPA1 signal in the surface of unpermealized T cells (Fig 1b). We probed surface as well as total expression of TRPA1 in murine T cells that are at resting (naïve) stage and/or activated with either the mitogen Concanavalin A (ConA, a lectin that acts as a mitogen and results in T cell activation) or by T cell Receptor (TCR) stimulation with α-CD3/α-CD28 antibodies [17, 18].

**Figure 1.**
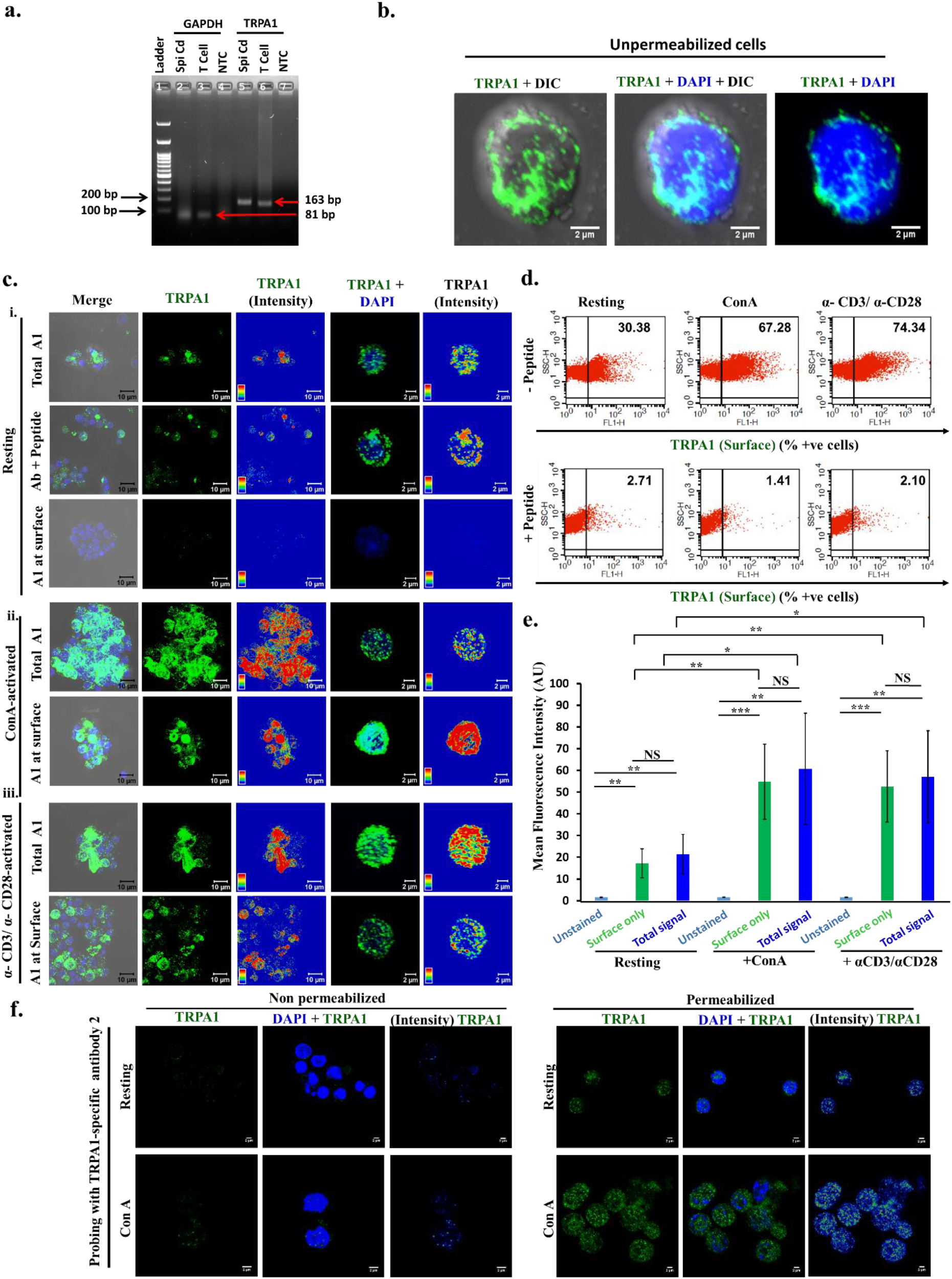
Endogenous expression of TRPA1 primary murine T cells. **a.** RT-PCR for TRPA1 and GAPDH from mRNA isolated from T cell. Total mRNA isolated from mouse spinal cord is used as a positive control and no-template control (NTC) is used as negative control. **b.** Immunolocalization of TRPA1 in the surface of unpermeabilized T cells. **c.** Expression of TRPA1 is increased in activated T cells compared to the resting conditions. TRPA1 present at the surface only (in non permeabilized cells) and in whole cell (in permeabilized cells) were probed. Triton X-100 permeabilized cells when stained with TRPA1 specific antibody pre-incubated with its antigenic peptide results in loss of TRPA1 signal, indicating the specificity of the antibody. Confocal images of purified CD3^+ve^ murine T cells stained with antibody detecting the extracellular loop of TRPA1 are shown. Both surface level and total expression of TRPA1 in resting cells (**i**), in ConA-activated cells (**ii**), and in α-CD3/α-CD28-activated T cells (**iii**) are shown. Fluorescence intensity of the TRPA1 signal is indicated in rainbow scale (right panels). **d.** Dot-plot values indicating the percentage of cells expressing TRPA1 at the surface. The number of TRPA1^+ve^ cells are much higher in activated conditions. This staining is completely blocked when the same antibody is pre-treated with its specific blocking peptide. **e.** Mean fluorescence intensity determined by Flow cytometry analysis reveal that ConA-activated and α-CD3/α-CD28-activated T cells have higher levels of TRPA1 than the Resting T cells. The difference in TRPA1 expression level (both in surface as well in whole cell) between resting stage with ConA-activated or α-CD3/α-CD28-activated T cells are significant. The p values are: ns = non-significant; * = < 0.05; ** = <0.01; *** = <0.001. **f.** Immunodetection of TRPA1 in T cell by using another antibody recognizing the epitope present in the N-terminal cytoplasmic domain. In permeabilized cells this antibody detects TRPA1 at low level in resting condition and modest level in ConA-treated condition. This antibody fails to detect TRPA1 in unpermeabilized T cells.

Further confocal microscopy analysis revealed that TRPA1 is endogenously expressed in resting and activated T cells as distinct clusters that are primarily located at the cell surface (Fig. 1c). Notably, the TRPA1 signal was blocked upon pre-incubating the antibodies with their antigenic peptide confirming the specificity of the antibody used (Fig. 1c). The intracellular localization of TRPA1 was almost minimal as there was no significant difference in its expression in surface versus in whole cell (in resting conditions). However, all the T cells do not express TRPA1 at resting state. Flow cytometry results suggest that the expression level of TRPA1 was increased in Con A activated and in TCR activated T cells (Fig. 1d & 1e).

To confirm the endogenous expression of TRPA1 in T cell, we used another antibody (Novus Biologicals) raised against epitope present in the N-terminal cytoplasmic domain of TRPA1. This antibody detects TRPA1 modestly in resting T cells and strongly in ConA-treated T cells, but after permeabilization (Fig. 1f). This antibody does not detect TRPA1 in unpermeabilized T cells. Taken together, the data strongly suggest that TRPA1 is endogenously expressed in murine T cells.

Similar to murine T cells, human PBMC-derived T cells also show TCR and ConA activation driven increased expression of TRPA1 (Fig. 2). The Z-section images showed increase in TRPA1 at the surface on T cell activation by ConA or TCR (Fig. 2). These qualitative and quantitative data strongly suggest for endogenous expression of TRPA1 in T cells and increased surface expression of TRPA1 correlates with T cell activation process.

**Figure 2.**
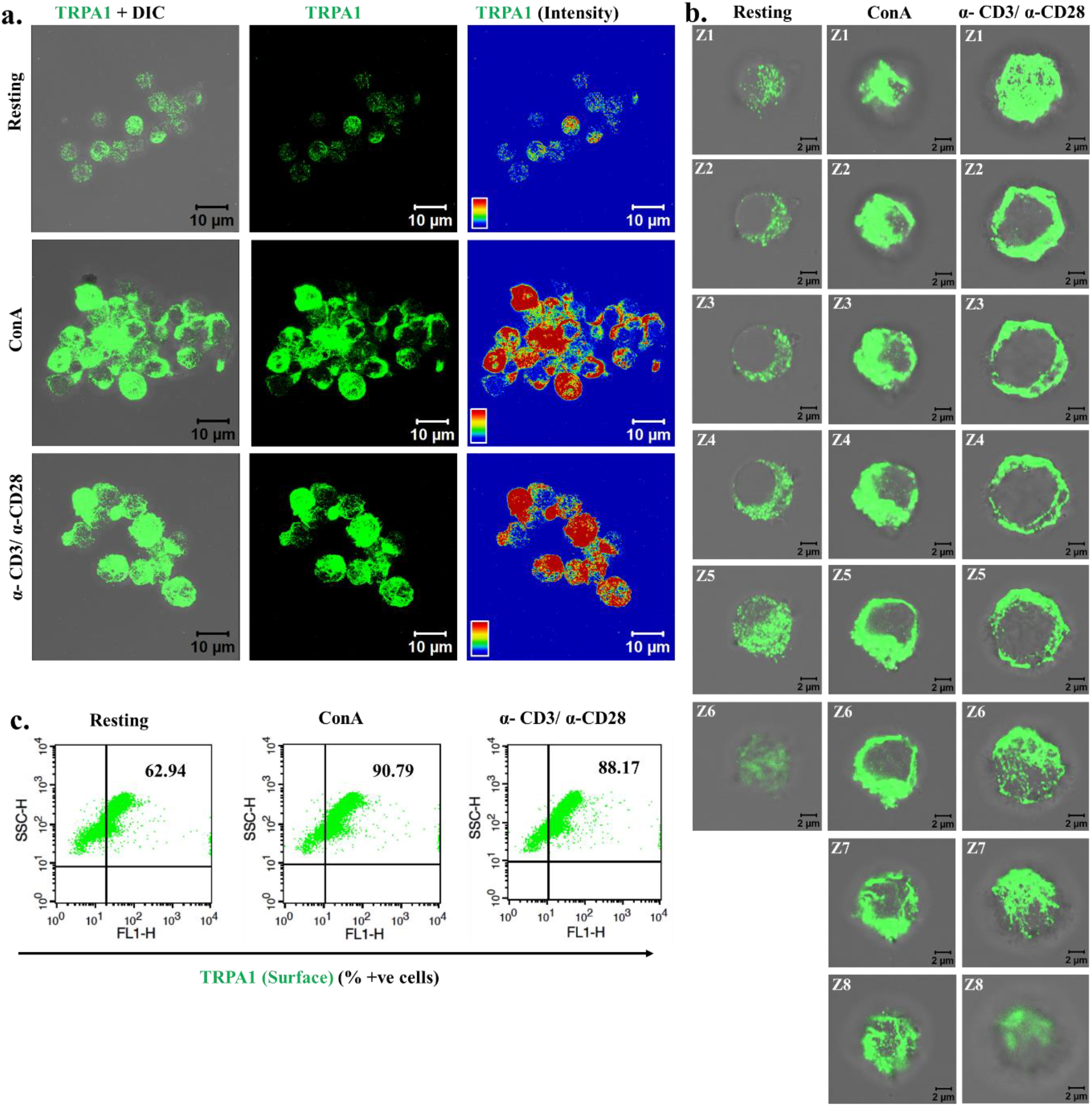
Human T cells show increased surface expression of TRPA1 upon activation. **a.** Confocal imaging revealed that TRPA1 is endogenously expressed in purified CD3^+ve^ human T cells and TRPA1 expression increases in activated T cells. **b.** Optical sections (Z1-Z8) of T cells reveal that TRPA1 is present mostly at or near the cell surface both in resting and in activated conditions. **c**. Flow cytometry dot-plot values indicate that around 60% of human T cells are TRPA1^+ve^ under resting conditions while ~90% cells are TRPA1^+ve^ in immunologically activated state.

### TRPA1 activation induces increased Ca^2+^ level in T cells

In order to explore if the TRPA1 present in T cells are functional, we performed Ca^2+^-imaging experiments (Fig. 3). For that purpose, we have used purified mouse T cells loaded with Fluo-4 AM. Live cell imaging revealed that in the absence of any stimuli, there is no increase in Fluo-4 intensity in the majority of the cells with respect to time. However, upon stimulation by TRPA1 activator AITC, intracellular Ca^2+^ level increase in most of the T cells (Fig 3a). This AITC-mediated increase in Ca^2+^-level was effectively blocked by TRPA1-specific inhibitors A-967079, and HC-030031 (Fig 3& 4). Calcium chelation experiments with BAPTA-AM as well as by EGTA (present in extracellular media) suggest that TRPA1 regulates Ca^2+^influx from both extracellular and intracellular reserves (Fig 4). Similarly, the level of intracellular Ca^2+^-goes down after adding the inhibitor in activator (AITC) treated cells (Fig 4f). To confirm all these effects in a more quantitatively, we compared the level of Ca^2+^ in T cells just before and after adding the TRPA1 modulatory drugs (Fig 4g). Such analysis clearly suggest the changes in the fluorescence intensity in quick time. This confirmed that functional TRPA1 is expressed in T cells.

**Figure 3.**
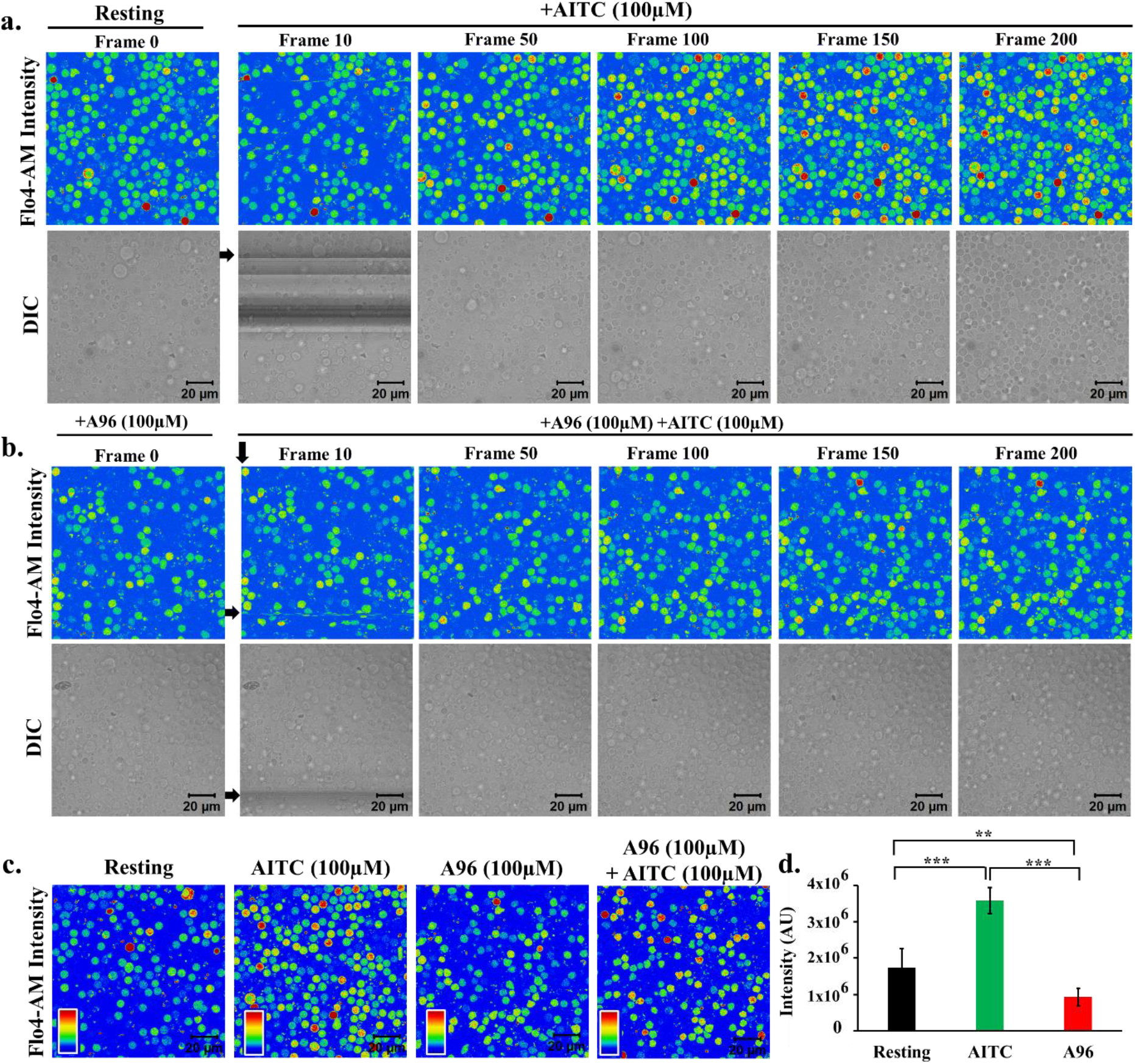
TRPA1 activation induces increase in intracellular Ca^2+^ levels in murine T cells. **a-b.** Fluorescence intensity images derived from time-series imaging of view fields containing multiple cells loaded with Ca^2+^-sensing dye Fluo4-AM are depicted here. The time difference between each frame is 5 seconds. The cells were treated with different pharmacological agents at the 10^th^ frame (F10). Activation of TRPA1 by its specific activator Allyl isothiocyanate (AITC 100uM) causes increment in the Ca^2+^-level (A) which can be blocked by pre-incubating the cells with TRPA1-specific inhibitor A967079 (A96; 100 μM) for 30 min (B). **c.** Fluo-4 intensity of T cells at resting stage or incubated with AITC (100μM), A96 (100μM) or combination of AITC (100μM) andA96 (100μM) for 12 hours is shown. **d.** Quantification of Fluo-4 intensity from 6 random fields after acquisition of 200 frames (1000 secs) of two independent experiments is depicted. AITC (100μM) treatment significantly increases Ca^2+^-levels, while A96 (100 μM) treatment reduces intracellular Ca^2+^-levels below that of resting T cells. The p values are: ns = non-significant; * = < 0.05; ** = <0.01; *** = <0.001.

**Figure 4:**
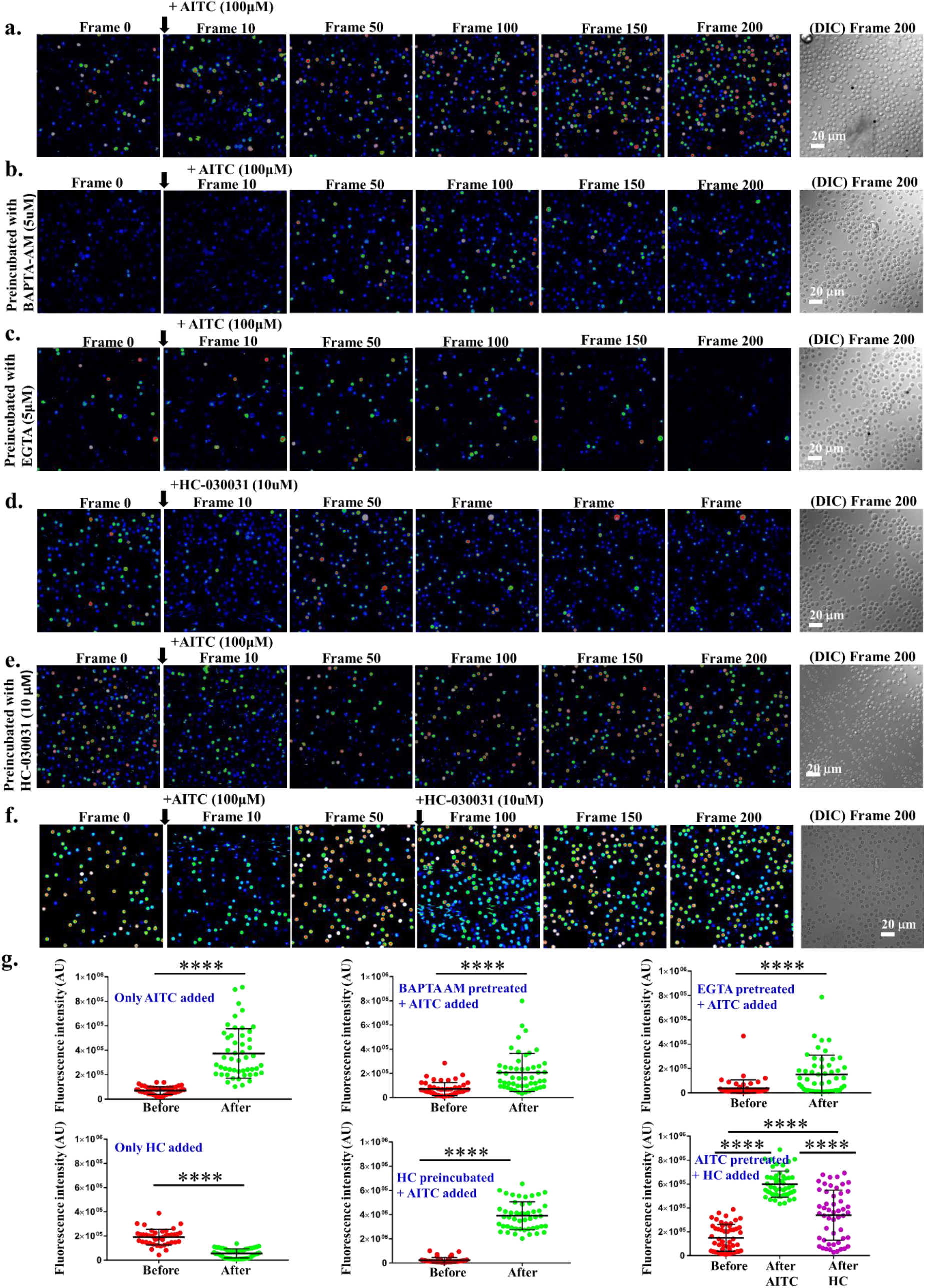
TRPA1 activation-induced increase in intracellular Ca^2+^ can be altered by Ca^2+^-chelators and channel inhibitors. Fluorescence intensity images derived from time-series imaging of view fields containing multiple cells loaded with Ca^2+^-sensing dye Fluo4-AM are depicted here. The time difference between each frame is 5 seconds. Untreated cells at resting stage or preincubated with specific Ca^2+^-chelators or TRPA1 blocker. Activation of TRPA1 by AITC at the 10^th^ frame causes increment in the Ca^2+^-level (a) which can be reduced by pre-incubating the cells with BAPTA-AM (intracellular Ca^2+^ chelator) (b) or with extracellular Ca^2+^ chelator (EGTA) (c). Application of HC-030031, another inhibitor of TRPA1 results in reduced intracellular level of Ca^2+^ (d). Application of AITC on population preincubated with HC-030031 fails to increase the basal Ca^2+^-levels (e). Application of HC-030031 after AITC results in initial rise (due to activation) followed by reduction (due to inhibition) of the basal Ca^2+^-levels (f). In each case, the DIC image represents the number and morphology of cells in the view-field at the end of the experiments (200^th^ frame). **g.** Quantification of basal Ca^2+^-levels in multiple cells before and after addition of TRPA1 modulatory agents. The maximum time gap between these two measurements are ~5 sec. **** indicates P value = ˂0.0001. For more details see supplementary movies.

### TRPA1 is involved in ConA/TCR-mediated T cell activation and effector cytokine secretion

In order to explore the role of TRPA1 in T cell activation, we have activated the cells either by ConA or via TCR stimulation and probed for the expression of activation markers, namely CD25 and CD69 in the purified murine T cell (CD90.2^+^) population. The expression of these markers was probed after ConA treatment with or without TRPA1 channel modulators (Fig. 5). Flow cytometric evaluation revealed a shift in the T cell population expressing CD25 upon T cell activation (in resting condition, 3.495 ± 0.95 %; in ConA-activated condition, 73.795 ± 0.82 %; in TCR -activated condition, 59.075 ± 1.14 %). Notably, in the presence of TRPA1 inhibitor (A-967079, 100µM), T cell activation by ConA or TCR was significantly inhibited (Fig. 5a, c). In that condition (after treatment with both ConA and A-967079), 27.87 ± 0.41 % of the cells express CD25. Down-regulation was seen when cells were treated with TCR along with A-967079, where only 30.495 ± 1.08 % cells are found to be CD25^+^.

**Figure 5.**
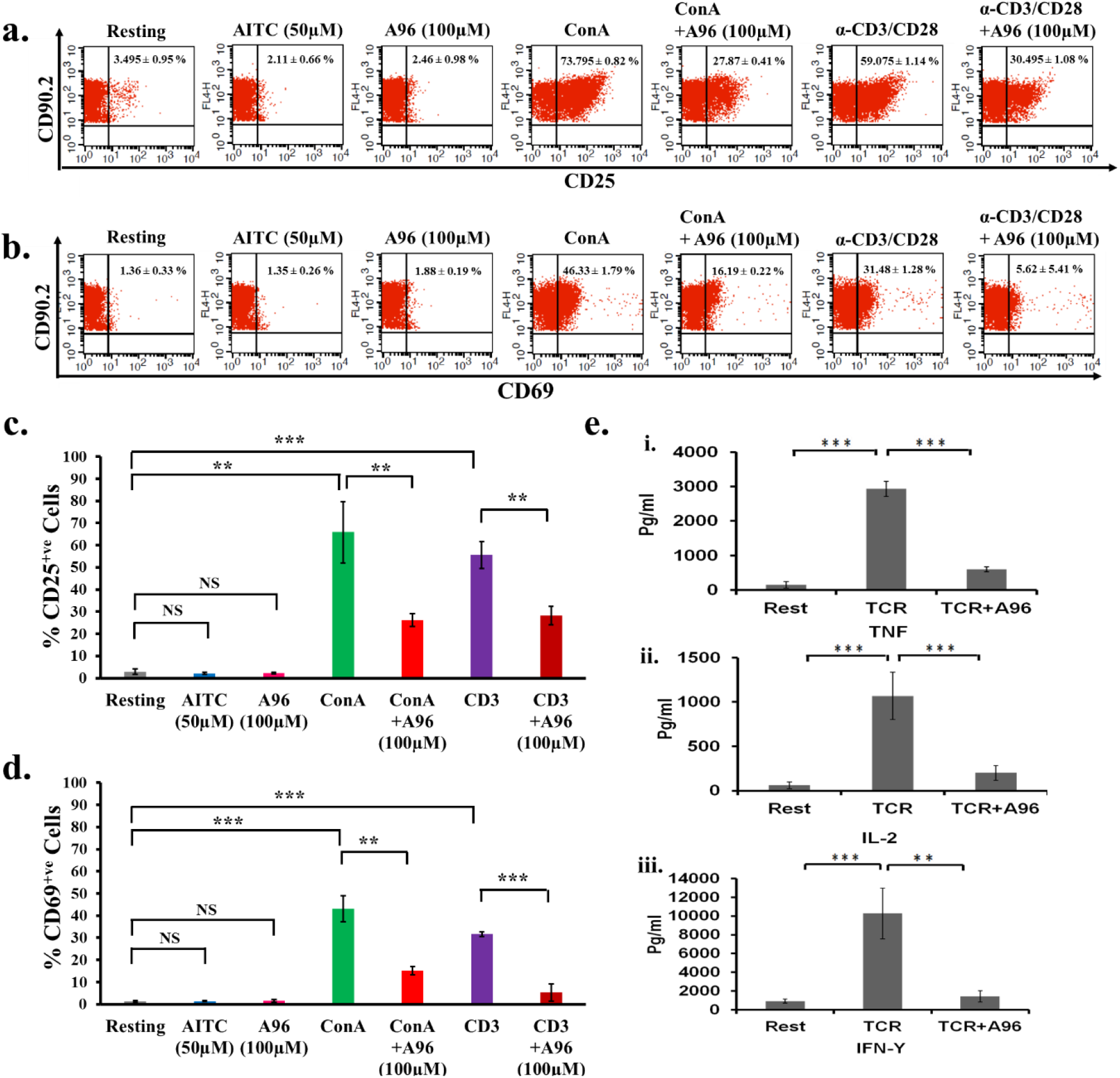
Pharmacological inhibition of endogenous TRPA1 blocks T cell activation. **a-b.** T cell activation markers CD25 and CD69 were analyzed by flow cytometry after incubating the cells with TRPA1 modulators for 36 hours. Inhibition of TRPA1 by A96 (100µM) reduces the ConA-mediated activation. Treatment of murine T cells with AITC (50µM) or A96 (100µM) alone do not increase the % of CD25^+^ cells or % of CD69^+^ cells. The average number ± SD values of CD25^+^ or CD69^+^ cells are mentioned in the upper right corner of each dot-plot. Representative dot plots of 3 independent experiments are shown. **c-d**. T cells treated with A96 (100µM) along with ConA (5 µg/ml) or plate-bound α-CD3 (2µg/ml) and soluble α-CD28 (2µg/ml) have reduced percentage of CD25^+^ or CD69^+^ cells. The corresponding levels/intensity of CD25 or CD69 expression (determined from MFI values) in response incubation with indicated modulators are shown. (n = 3). **e.** Graphical bars represent the concentration (in pg/ml) of effector cytokines TNF (**i**), IL-2 (**ii**), IFNγ (**iii**) released from T cells around 36 hours. Incubation of TRPA1 inhibitor A96 (100µM) along with plate-bound α-CD3 (2µg/ml) and soluble α-CD28 (2µg/ml) resulted in significant reduction in the release of cytokines TNF, IL-2 and IFNγ. The p values are: ns = non-significant; ** = <0.01; *** = <0.001. (n = 3 independent experiments).

In a similar manner, TRPA1 inhibitor (A-967079) also reduced CD69 expression. The effect of TRPA1 inhibitor on CD69 expression was also reflected by the percentage of CD69 positive cells (in resting condition, 1.36±0.33%; in ConA-activated condition, 46.33±1.79%; in TCR-activated 31.48±1.28%) which was reduced after treating with the inhibitor (in combination of ConA and A-967079, 16.19 ± 0.22 %; in TCR and A-967079 combination, 5.62 ± 5.41 %) (Fig. 5b, 5d).

T cell activation involves an increased level of secretion of several effector cytokines like TNF, IFNγ, and IL-2. We explored the role of TRPA1 in the production of these cytokines by analyzing the culture supernatants by ELISA (Fig. 5e). TCR stimulation induced high levels of TNF (Resting condition: 151.8±94.88 pg/ml; TCR-mediated activated condition: 2931.98±220.85 pg/ml) while inhibition of TRPA1 by A-967079 blocked the effect of these stimulators on TNF production (TCR + A-967079 combination: 597.74±71.83 pg/ml) (Fig. 5e). TRPA1 inhibition also blocked IL-2 secretion (Resting: 62±37.91 pg/ml; TCR: 1067.4 ± 267.003 pg/ml; TCR + A-967079 combination: 200.33±83.25 pg/ml) (Fig. 5e). Moreover IFNγ production was down-regulated by A-967079 treatment (Resting: 912.83±216.20pg/ml; TCR: 10279.5 ± 2702.65 pg/ml; TCR + A-967079 combination: 1421.72±599.3 pg/ml) (Fig. 5e). Taken together the results indicate that TRPA1 is expressed in T cells and may positively regulate T cell activation and effector cytokine release.

## Discussion

T cell activation involves several distinct signaling events, and the influx of Ca^2+^-ions is vital during this process. However, so far only a few Ca^2+^ channels have been detected in T cells, whereas the identity of the major Ca^2+^ channels present in T cells is unknown. In addition, the mode of regulation of these channels and their exact role in the context of T cell functions are mostly unknown. TRPA1 can be activated by several means and also by endogenous factors. Though TRPA1 has been considered as a “cold-activated ion channel,” it’s true thermosensitive nature is debatable. For example, it has been shown that TRPA1 is not directly gated by cold but rather gated by increased intracellular Ca^2+^-levels as a consequence of cooling [19]. It is important to mention that Ca^2+^ ions directly gate hTRPA1 in a PLC-independent fashion. This is confirmed by the fact that TRPA1 shows Ca^2+^-induced currents even after treatment with the PLC-blocker U73122 (10 µM) [3].

In this work, we demonstrated that functional TRPA1, a non-selective cation channel is expressed in the human and murine T cells. Our results suggest that TRPA1 plays an important role relevant for T cell activation, likely by different endogenous factors. In a recent report, the functional expression of TRPA1 in murine and human CD4^+^ T cells has been shown to regulate T cell activation and pro-inflammatory responses [20]. However, the previous reports have suggested a pro-inflammatory role of TRPA1 towards immune activation and inflammation [21]. So far, TRPA1 has been considered as one of the critical regulators of neurogenic inflammation and neuropeptide release [22]. It has also been reported that TRPA1 is associated with inflammation and puritogen responses in dermatitis [23]. In fact expression of TRPA1 in the peripheral sensory neurons and its involvement in the neural excitation has been demonstrated. However, limited information is available currently about the expression and role of TRPA1 in non-neuronal cells. The association of TRP channels towards inflammation and immunogenic responses has been largely found to be positively regulated during the immune-physiology of the cellular and systemic responses. TRPA1 has been reported to contribute to the inflammation induced pain and associated experimental colitis in mice models [23–26]. Interestingly, recently it has been reported that CD4^+^ T cell-associated TRPA1 may regulate inflammation in experimental colitis model in mice [20]. However, evidence for any such functional requirement of TRPA1 in T cell responses, *in vitro*, remains scanty.

In this work, the endogenous expression of TRPA1 in primary murine and human T cell was observed by using two different TRPA1 specific antibodies. Expression of TRPA1 was found to be mostly predominant at the surface of these cells rather than being present in intracellular regions. We also provided evidence for the functional role of TRPA1 in immune functions. Flow cytometric analysis coupled with confocal imaging conclusively suggested enhanced expression of TRPA1 (both at surface level and in total) in activated T cells as compared to resting conditions. TRPA1 activation by specific ligands leads to increased intracellular Ca^2+^ concentration in purified murine T cells, confirming that this channel is present in a functional form in resting T cells. Inhibition of TRPA1 by its specific inhibitor reduces TCR and Con A-driven mitogenic activation of T cells. This suggests that TRPA1 might be involved in the signaling pathways towards T cell activation and thus inhibiting TRPA1 activity has profound inhibitory effects on CD25 and CD69 expression together with the secretion of signature effector cytokines such as TNF, IFN-γ, and IL-2. TNF production is associated with several pro-inflammatory responses [27]. Additionally TNF is shown to be a major mediator of different inflammatory disease conditions like colitis and rheumatoid arthritis (RA) [28, 29]. TRPA1 channel expression is found to be increased in peripheral blood leukocytes of RA patients and associated with pain [30]. Moreover, it has been shown that TRPA1 facilitates TNF directed inflammatory responses in various pathophysiological conditions and blockade of TRPA1 receptors may be beneficial in reducing TNF induced the chronic pain [31, 32]. Interestingly, it has been shown that TRPA1^−/−^T cells may produce higher IL-2 and IFN-γ but not TNF during TCR stimulation [20]. Functional role of IFN-γ as a signature Th1 cytokine is also implicated in pro-inflammatory responses and disease conditions like colitis [33]. However, there are some reports which suggest an anti-inflammatory role of IFN-γ in the mouse model of colitis [34, 35]. Apparently, these reports indicate that these cytokines may differentially regulate inflammatory responses in different disease conditions. Interestingly, we have found that during *in vitro* TCR activation the induction of signature Th1 and pro-inflammatory cytokines like IL-2, IFN-γ, and TNF could be down-regulated by the TRPA1 specific inhibitor, A-967079. Moreover, ConA and TCR stimulated induction of T cell activation markers like CD25 and CD69 were also found to be downregulated in presence A-967079. TRPA1 upon activation by its specific activator (AITC) was found to increase the Ca^2+^-levels, while two different inhibitors namely A-967079 and HC-030031 are able to reduce the intracellular Ca^2+^-levels. Therefore, it seems that during T cell activation, TRPA1 becomes functional and shows augmented expression and executes immune-regulatory functions, whereas, inhibition of this channel inhibits T cell activation. Recent report demonstrating the activation of TRPA1 by specific microRNA can also be relevant for T cell activation [36]. While our current work confirms the functional expression of TRPA1 channel and its requirement towards T cell activation, the involvement of this ion channel in T cell activation process associated with inflammatory diseases needs further investigation.

In brief, our current findings demonstrated an *in vitro* T cell activation directed functional expression and requirement of TRPA1 in T cells. This is in line with the earlier reports of inflammatory responses associated to the function of TRPA1 in various physiological systems. The current observation might have implication in the immunogenic and inflammatory role of T cell responses as well.

## List of abbreviations

AITC: Allyl isothiocyanate
BAPTA-AM: 1,2-Bis(2-aminophenoxy)ethane-N,N,N′,N′-tetraacetic acid tetrakis (acetoxymethyl ester)
CD25: Cluster of Differentiation 25
CD3^+^: Cluster of Differentiation 3
CD69: Cluster of Differentiation 69
ConA: ConcanavalinA
DCs: Dendritic Cells
ELISA: Enzyme Linked Immunosorbent Assay
EGTA: Ethylene glycol-bis(β-aminoethyl ether)-*N*,*N*,*N*′,*N*′-tetraacetic acid
HNE: 4-hydroxy-2-nonenal
hPBMC: Human Peripheral Blood Mononuclear Cells
IFNγ: Interferon-γ
IL-2: Interleukin-2
LPS: Lipopolysaccharide
NO: Nitric Oxide
PKC: Protein Kinase C
PLC: Phospholipase C
ROS: Reactive Oxygen Species
RT-PCR: Real Time Polymerase Chain Reaction
TNF: Tumor Necrosis Factor
TRPV: Transient Receptor Potential cation channel subfamily Vanilloid
TRPA1: Transient Receptor Potential Ankyrin1
Th1: T helper type1
Th2: T helper type2

## Acknowledgment

We gratefully acknowledge the support from the Imaging Facility, Flow Cytometry Facility of NISER and the Animal house facilities of NISER and ILS, Bhubaneswar.

## Data availability

All data generated during this study are included in this published article.

## Author contributions

SSS, RKM, CG, and SC conceived the idea and designed all the experiments. SSS, RKM, TA, AT, SS, and PSK performed all the experiments. SSS, RKM, AT, TA, SS, PSK, CG, and SC analyzed the data. SSS, RKM, CG, and SC wrote the manuscript. CG communicated the manuscript.

## References

[1] L.J. Wu, T.B. Sweet, D.E. Clapham, International Union of Basic and Clinical Pharmacology. LXXVI. Current progress in the mammalian TRP ion channel family, Pharmacol Rev 62(3) (2010) 381–404.

[2] G.M. Story, A.M. Peier, A.J. Reeve, S.R. Eid, J. Mosbacher, T.R. Hricik, T.J. Earley, A.C. Hergarden, D.A. Andersson, S.W. Hwang, P. McIntyre, T. Jegla, S. Bevan, A. Patapoutian, ANKTM1, a TRP-like channel expressed in nociceptive neurons, is activated by cold temperatures, Cell 112(6) (2003) 819–29.

[3] J.F. Doerner, G. Gisselmann, H. Hatt, C.H. Wetzel, Transient receptor potential channel A1 is directly gated by calcium ions, J Biol Chem 282(18) (2007) 13180–9.

[4] K.Y. Kwan, D.P. Corey, Burning cold: involvement of TRPA1 in noxious cold sensation, J Gen Physiol 133(3) (2009) 251–6.

[5] M. Bandell, G.M. Story, S.W. Hwang, V. Viswanath, S.R. Eid, M.J. Petrus, T.J. Earley, A. Patapoutian, Noxious cold ion channel TRPA1 is activated by pungent compounds and bradykinin, Neuron 41(6) (2004) 849–57.

[6] S. Feske, Calcium signalling in lymphocyte activation and disease, Nat Rev Immunol 7(9) (2007) 690–702.

[7] M. Vig, J.P. Kinet, Calcium signaling in immune cells, Nat Immunol 10(1) (2009) 21–7.

[8] Z. Petho, A. Balajthy, A. Bartok, K. Bene, S. Somodi, O. Szilagyi, E. Rajnavolgyi, G. Panyi, Z. Varga, The anti-proliferative effect of cation channel blockers in T lymphocytes depends on the strength of mitogenic stimulation, Immunol Lett 171 (2016) 60–9.

[9] M. Trevisani, J. Siemens, S. Materazzi, D.M. Bautista, R. Nassini, B. Campi, N. Imamachi, E. Andre, R. Patacchini, G.S. Cottrell, R. Gatti, A.I. Basbaum, N.W. Bunnett, D. Julius, P. Geppetti, 4-Hydroxynonenal, an endogenous aldehyde, causes pain and neurogenic inflammation through activation of the irritant receptor TRPA1, Proc Natl Acad Sci U S A 104(33) (2007) 13519–24.

[10] L. Cruz-Orengo, A. Dhaka, R.J. Heuermann, T.J. Young, M.C. Montana, E.J. Cavanaugh, D. Kim, G.M. Story, Cutaneous nociception evoked by 15-delta PGJ2 via activation of ion channel TRPA1, Mol Pain 4 (2008) 30.

[11] T.E. Taylor-Clark, S. Ghatta, W. Bettner, B.J. Undem, Nitrooleic acid, an endogenous product of nitrative stress, activates nociceptive sensory nerves via the direct activation of TRPA1, Mol Pharmacol 75(4) (2009) 820–9.

[12] F. Cevikbas, X. Wang, T. Akiyama, C. Kempkes, T. Savinko, A. Antal, G. Kukova, T. Buhl, A. Ikoma, J. Buddenkotte, V. Soumelis, M. Feld, H. Alenius, S.R. Dillon, E. Carstens, B. Homey, A. Basbaum, M. Steinhoff, A sensory neuron-expressed IL-31 receptor mediates T helper cell-dependent itch: Involvement of TRPV1 and TRPA1, J Allergy Clin Immunol 133(2) (2014) 448–60.

[13] V. Meseguer, Y.A. Alpizar, E. Luis, S. Tajada, B. Denlinger, O. Fajardo, J.A. Manenschijn, C. Fernandez-Pena, A. Talavera, T. Kichko, B. Navia, A. Sanchez, R. Senaris, P. Reeh, M.T. Perez-Garcia, J.R. Lopez-Lopez, T. Voets, C. Belmonte, K. Talavera, F. Viana, TRPA1 channels mediate acute neurogenic inflammation and pain produced by bacterial endotoxins, Nat Commun 5 (2014) 3125.

[14] R.K. Majhi, S.S. Sahoo, M. Yadav, B.M. Pratheek, S. Chattopadhyay, C. Goswami, Functional expression of TRPV channels in T cells and their implications in immune regulation, FEBS J 282(14) (2015) 2661–81.

[15] S. Chattopadhyay, J. O’Rourke, R.E. Cone, Implication for the CD94/NKG2A-Qa-1 system in the generation and function of ocular-induced splenic CD8+ regulatory T cells, Int Immunol 20(4) (2008) 509–16.

[16] C. Goswami, H. Schmidt, F. Hucho, TRPV1 at nerve endings regulates growth cone morphology and movement through cytoskeleton reorganization, FEBS J 274(3) (2007) 760–72.

[17] B.L. Levine, J.D. Mosca, J.L. Riley, R.G. Carroll, M.T. Vahey, L.L. Jagodzinski, K.F. Wagner, D.L. Mayers, D.S. Burke, O.S. Weislow, D.C. St Louis, C.H. June, Antiviral effect and ex vivo CD4+ T cell proliferation in HIV-positive patients as a result of CD28 costimulation, Science 272(5270) (1996) 1939–43.

[18] J.E. Kay, Mechanisms of T lymphocyte activation, Immunol Lett 29(1-2) (1991) 51–4.

[19] S. Zurborg, B. Yurgionas, J.A. Jira, O. Caspani, P.A. Heppenstall, Direct activation of the ion channel TRPA1 by Ca^2+^, Nat Neurosci 10(3) (2007) 277–9.

[20] S. Bertin, Y. Aoki-Nonaka, J. Lee, P.R. de Jong, P. Kim, T. Han, T. Yu, K. To, N. Takahashi, B.S. Boland, J.T. Chang, S.B. Ho, S. Herdman, M. Corr, A. Franco, S. Sharma, H. Dong, A.N. Akopian, E. Raz, The TRPA1 ion channel is expressed in CD4+ T cells and restrains T-cell-mediated colitis through inhibition of TRPV1, Gut (2016).

[21] O. Gouin, K. L’Herondelle, N. Lebonvallet, C. Le Gall-Ianotto, M. Sakka, V. Buhé, E. Plée-Gautier, J.-L. Carré, L. Lefeuvre, L. Misery, R. Le Garrec, TRPV1 and TRPA1 in cutaneous neurogenic and chronic inflammation: pro-inflammatory response induced by their activation and their sensitization, Protein & Cell (2017) 1–18.

[22] D.M. Bautista, M. Pellegrino, M. Tsunozaki, TRPA1: A gatekeeper for inflammation, Annu Rev Physiol 75 (2013) 181–200.

[23] B. Liu, J. Escalera, S. Balakrishna, L. Fan, A.I. Caceres, E. Robinson, A. Sui, M.C. McKay, M.A. McAlexander, C.A. Herrick, S.E. Jordt, TRPA1 controls inflammation and pruritogen responses in allergic contact dermatitis, FASEB J 27(9) (2013) 3549–63.

[24] M. Miampamba, S. Chery-Croze, S. Detolle-Sarbach, D. Guez, J.A. Chayvialle, Antinociceptive effects of oral clonidine and S12813-4 in acute colon inflammation in rats, Eur J Pharmacol 308(3) (1996) 251–9.

[25] M.A. Engel, C. Becker, P.W. Reeh, M.F. Neurath, Role of sensory neurons in colitis: increasing evidence for a neuroimmune link in the gut, Inflamm Bowel Dis 17(4) (2011) 1030–3.

[26] D.M. Bautista, S.E. Jordt, T. Nikai, P.R. Tsuruda, A.J. Read, J. Poblete, E.N. Yamoah, A.I. Basbaum, D. Julius, TRPA1 mediates the inflammatory actions of environmental irritants and proalgesic agents, Cell 124(6) (2006) 1269–82.

[27] C. Popa, M.G. Netea, P.L. van Riel, J.W. van der Meer, A.F. Stalenhoef, The role of TNF-alpha in chronic inflammatory conditions, intermediary metabolism, and cardiovascular risk, J Lipid Res 48(4) (2007) 751–62.

[28] B.K. Popivanova, K. Kitamura, Y. Wu, T. Kondo, T. Kagaya, S. Kaneko, M. Oshima, C. Fujii, N. Mukaida, Blocking TNF-alpha in mice reduces colorectal carcinogenesis associated with chronic colitis, J Clin Invest 118(2) (2008) 560–70.

[29] H. Matsuno, K. Yudoh, R. Katayama, F. Nakazawa, M. Uzuki, T. Sawai, T. Yonezawa, Y. Saeki, G.S. Panayi, C. Pitzalis, T. Kimura, The role of TNF-alpha in the pathogenesis of inflammation and joint destruction in rheumatoid arthritis (RA): a study using a human RA/SCID mouse chimera, Rheumatology (Oxford) 41(3) (2002) 329–37.

[30] I. Pereira, S.J. Mendes, D. Pereira, T.F. Muniz, V.L. Colares, C.R. Monteiro, M.M.d.S. Martins, M.A. Grisotto, V. Monteiro-Neto, S.G. Monteiro, Transient receptor potential ankyrin 1 channel expression on peripheral blood leukocytes from rheumatoid arthritic patients and correlation with pain and disability, Frontiers in pharmacology 8 (2017) 53.

[31] E.S. Fernandes, F.A. Russell, D. Spina, J.J. McDougall, R. Graepel, C. Gentry, A.A. Staniland, D.M. Mountford, J.E. Keeble, M. Malcangio, S. Bevan, S.D. Brain, A distinct role for transient receptor potential ankyrin 1, in addition to transient receptor potential vanilloid 1, in tumor necrosis factor alpha-induced inflammatory hyperalgesia and Freund’s complete adjuvant-induced monarthritis, Arthritis Rheum 63(3) (2011) 819–29.

[32] A. Koivisto, H. Chapman, N. Jalava, T. Korjamo, M. Saarnilehto, K. Lindstedt, A. Pertovaara, TRPA1: a transducer and amplifier of pain and inflammation, Basic Clin Pharmacol Toxicol 114(1) (2014) 50–5.

[33] R. Ito, M. Shin-Ya, T. Kishida, A. Urano, R. Takada, J. Sakagami, J. Imanishi, M. Kita, Y. Ueda, Y. Iwakura, K. Kataoka, T. Okanoue, O. Mazda, Interferon-gamma is causatively involved in experimental inflammatory bowel disease in mice, Clin Exp Immunol 146(2) (2006) 330–8.

[34] S.Z. Sheikh, K. Matsuoka, T. Kobayashi, F. Li, T. Rubinas, S.E. Plevy, Cutting edge: IFN-gamma is a negative regulator of IL-23 in murine macrophages and experimental colitis, J Immunol 184(8) (2010) 4069–73.

[35] Y. Jin, Y. Lin, L. Lin, C. Zheng, IL-17/IFN-gamma interactions regulate intestinal inflammation in TNBS-induced acute colitis, J Interferon Cytokine Res 32(11) (2012) 548–56.

[36] T.D. Sheahan, J. Hachisuka, S.E. Ross, Small RNAs, but Sizable Itch: TRPA1 Activation by an Extracellular MicroRNA. Neuron 99(3) (2018) 421–422.

